# Light-controlled phosphorylation in the TrkA-Y785 site by photosensitive UAAs activates the MAPK/ERK signaling pathway

**DOI:** 10.1101/2022.01.24.477628

**Authors:** Shu Zhao, Dong Liu

## Abstract

TrkA is a membrane receptor that upon ligand binding, induces autophosphorylation of tyrosine residues in the intracellular domain, and this process includes sites in the kinase activation loop (Y670, Y674, Y675) and two direct sites involved in downstream signaling pathways (Y490, Y785). At present, researchers cannot fully elucidate the regulatory mechanism of TrkA phosphorylation because TrkA signaling is a highly dynamic process, and a strategy with high temporal and spatial resolution will be beneficial to the mechanism research. Our previous study proposed a design scheme for photosensitive TrkA, which utilizes a new molecular light control technology to target TrkA-Y490 and three kinase domain sites (Y670, Y674, and Y675) through Genetic Code Expansion (GCE) technology combined with site-directed mutagenesis. We chose two light-controllable unnatural amino acids (UAAs) to introduce at the specific phosphorylation sites of the target protein TrkA. We focused on the regulation mechanism of these sites on the MAPK/ERK pathway downstream of TrkA. However, this method has not yet been validated for the TrkA-Y785 site. Therefore, this paper will continue to test the light-controlled method we established earlier in the Y785 site. We aim to improve further the experimental model of light-controlled phosphorylation of TrkA that we have established and finally lay the foundation for the comprehensive analysis of kinase-related pathways.

## Introduction

Tropomyosin receptor kinase A (TrkA) is a member of the Receptor Tyrosine Kinase (RTK) superfamily. It is widely involved in the cell physiological process and has significant effects on cell differentiation, growth, and synaptic plasticity. TrkA signaling pathways are closely related to cell growth, differentiation, and proliferation^1^. There are five crucial tyrosine (Tyr) residues in the cytoplasm of TrkA. Among them, Y670, Y674, Y675 are located in the kinase domain region, which are closely related to the activation of the enzyme^2–3^. Y490 and Y785 are the two main sites for activating TrkA downstream related signaling pathways outside the kinase domain region. The Y490 site is associated with Shc and Src domains to further start the Raf/MEK/ERK signaling pathway^4^, while the Y785 site mainly activates the PLCγ pathway^5^. The general concept of TrkA activation is that when neurotrophic factors (mainly NGF) stimulate TrkA, it will cause the autophosphorylation of tyrosine residues in the intracellular domain and then recruit the corresponding effector proteins to activate downstream signaling pathways. This activation effect is thought to be related to the five phosphorylation sites (Y490, Y670, Y674, Y675, Y785) and their combinations in the intracellular domain of TrkA, and the mechanism of TrkA phosphorylation has not been fully elucidated yet.

To better analyze the TrkA signaling pathway, we made full use of the latest advances in applying genetic code technology in mammals in our previous studies^6,7,8^. Our strategy was to regulate the phosphorylation state of the tyrosine side chain by introducing photosensitive UAAs (Unnatural Amino Acids, UAAs), thus making the phosphorylation of TrkA artificially controllable. We chose two developed photosensitive tyrosine analogs: 1) caged tyrosine (ONB)^9,10^, which can revert to tyrosine upon release of the photolabile moiety from the side chain upon light exposure; 2) azido-L-phenylalanine (AzF)^11^, which can crosslink with nearby molecules within a distance of 3–4 Å after light stimulation. The encoding of AzF and ONB into protein was to use genetic code expansion technology to suppress the expression of the stop codon (amber) in the mRNA of the target gene (here, TrkA) with the help of two pairs of engineered orthogonal aminoacyl tRNA synthetase (aaRS)/suppressor tRNA. The amber codon was used to encode a specific UAA, thereby enabling the read-through of the site-specific UAAs in the protein of interest^12,13^.

At present, we have successfully used mammalian cell models (PC12, HEK293T, and SH-SY5Y cells) to introduce AzF and ONB at the four phosphorylation sites (Y490, Y670, Y674, Y675) of TrkA^14^, respectively. And when these two photosensitive molecules were introduced into these sites, they all showed sensitivity to light and can specifically activate the downstream MAPK/ERK signaling pathway under the drive of U.V. stimulation. As one of the main docking sites for TrkA, Y785 has been reported in previous studies to act on the ERK pathway depending on PLCγ and PKC protein, and it can regulate the differentiation of PC12 cells together with TrkA-Y490^15,16^. But this light-controlled method we established before has not been tested in the Y785 site. We speculate that after the introduction of ONB at the TrkA-Y785 site, the downstream MAPK/ERK signaling pathway can also be activated by the light-controlled test according to the previous study, while the introduction effect of AzF at Y785 is unknown. It is expected that applying this light-controlled strategy to the TrkA-Y785 site will help us further study the regulatory mechanism of TrkA phosphorylation. Therefore, this paper intends to express the two photosensitive UAAs at the TrkA-Y785 site based on our previous research and combine the light-controlled method to detect its roles in regulating the MAPK/ERK signaling pathway. You can see the graphical Abstract in figure 1. Our research will expand the application system of genetic code expansion technology (GCE) in mammalian cells and provide a basis for understanding the contribution of the five phosphorylation sites of TrkA to specific signaling pathways (MAPK/ERK pathway).

**Figure 1.**
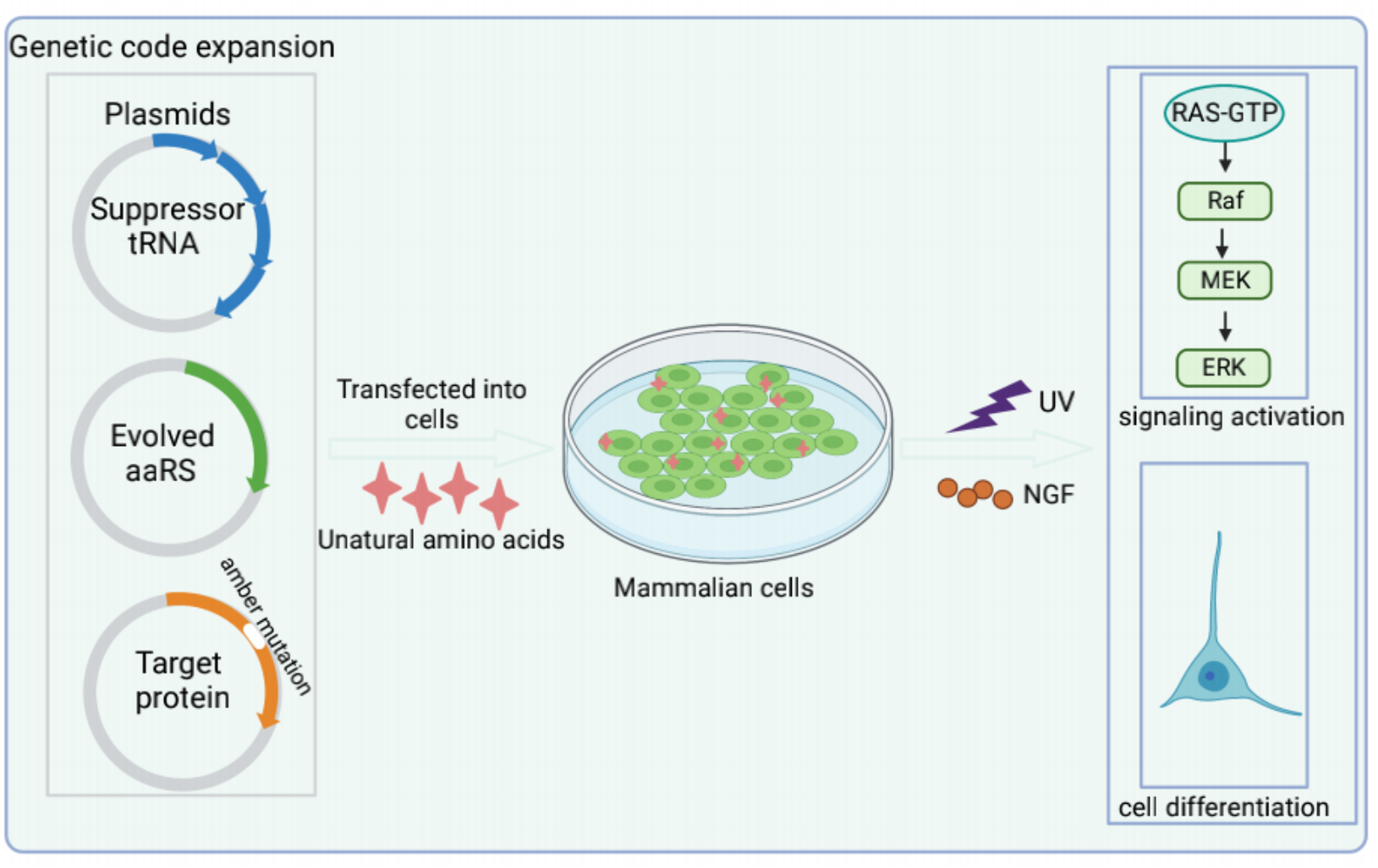
Graphical Abstract. Using gene encoding technology and light control method to express light-sensitive protein mutants in mammalian cells and dynamically regulate the activity of downstream signaling pathways and cell growth status.

## Materials and methods

### Plasmids and Cell Lines

The cell lines HEK293T, PC12, SH-SY5Y required for this experiment were purchased from ATCC; the TrkA-Y785 point mutation plasmid was constructed according to the single site-directed mutagenesis kit; The wild-type TrkA plasmid (PLNCX-TrkA-ΔO-GFP) **was** from Brain Lab (UPMC, Paris). For AzF, ONB, two orthogonal R.S./tRNA pairs that specifically recognize and fit them were required.

The plasmid AzFRS-(tRNA)_4X_^17^, which recognizes AzF, includes AzFRS (TyrRS mutant derived from E. coli) and amber stop codon tRNA (Tyr derived from Bacillus stearothermophilus). The plasmid was obtained from Dali Li lab (ECNU). The plasmid pONBYRS/U6-PyltRNA, which recognizes “photo-caged” tyrosine ONB, carries the gene pONBYRS derived from a M. pasteurii pyrRS mutant the stop codon tRNA comes from the PylRS/tRNA_CUA_ pair, which has also been reported previously^18^. This plasmid was obtained from Peng Cheng lab (Tsinghua).

### The expression of TrkA-785AzF, 785ONB mutants in HEK293T, SH-SY5Y cells

The TrkA-Y785amb plasmids in the experiments were co-transfected with R.S., tRNA (R.S./tRNA pairs corresponding to AzF or ONB) into cells. The experimental procedures are as follows: (1) Preparation before transfection: cells were plated the day before transfection to ensure that the cell density of the experimental group reached about 70% the next day. (2) One hour before transfection, changed medium to DMEM/high glucose medium (serum-free). (3) Preparation of PEI-DNA mixture: Take a 24-well plate as an example, mix 0.5 ug AzFRS-(tRNA)_4X_ plasmid with 0.5 ug PLNCX-TrkA (Y785amber)-ΔO-GFP plasmid (six-well plate DNA dosage is 2 ug) Cotransfected the mixture in 50 ul empty DMEM medium, marked as “A”; DNA (ug): PEI (ul) = 1:3, 3 ul PEI was added to another centrifuge tube containing 50 ul empty DMEM medium, marked as “B”; after standing for 5 min, add “B” to “A”. (4) After 20 min, drop the “A.B.” mixture into the corresponding well plate. (5) After 4-6 h, change the medium to the corresponding DMEM medium containing 1 mM AzF (for expressing ONB, replace it with fresh medium containing 100 uM ONB). Be careful to avoid light. (6) The cells were collected 24-48 h after transfection for fluorescence observation or western blot analysis. Suppression of the stop codon in TrkA-Y785amb produces fulllength EGFP whose fluorescent signal reports the binding of AzF or ONB at the stop codon TAG. 1 mM AzF or 100 um ONB was added to the cell growth medium, and the fluorescence expression was observed at 24-48 h. If the fluorescence intensity of EGFP in the cells was significantly enhanced compared with that without AzF or ONB, it indicated that AzF or ONB was successfully added to TrkA.

### Analysis of ONB and AzF incorporation efficiency at Y785 site of TrkA in HEK293T cells by fluorescence microscopy

24-48 h after transfection, photos were taken using a fluorescence microscope, and 6 different cell fields were taken for each mutant for statistics. The experiment was repeated no less than 3 times for each mutant. The pictures taken were processed by the software imageJ, and the average fluorescence intensity was counted. The rates of ONB and AzF incorporation at the Y785 site were expressed as the mean fluorescence intensity of cells incorporating ONB or AzF divided by the mean fluorescence intensity of cells expressing WT TrkA. Use the R language to draw statistical graphs.

### NGF and U.V. treatment for protein expression and cell phenotype analysis

48 h after transfection, NGF stimulation, and light treatment were performed. Before this step, the medium supplemented with 1 mM AzF or 100 um ONB was replaced with fresh PBS buffer. Then place the cell plates that need to be exposed to U.V. light on ice and irradiate them with light for 5-10 min in a 365 nm U.V. cross-linker (40 W). After U.V. treatment, the PBS was replaced with serum-free medium, and cells were cultured for 1 h in a carbon dioxide (5%) cell incubator at 37 °C. cells were then stimulated by adding human NGF to the medium at a 50 ng/ml final concentration. As a control, use an equal volume of NGF-free medium-well plates. HEK293T cells needed to be incubated at 37°C for 20 min for Western Blot analysis, while neuronal cells needed to be treated in NGF-containing medium for 24-48 h (dark) for cell growth and differentiation ratio analysis (No NGF treatment group also needed to ensure the same treatment time).

### Detecting the expression levels of TrkA protein and its activated downstream signaling proteins in HEK293T cells by western blot

Collected HEK293T cells 48 hours after transfection, washed with PBS, and prepared protein samples; quantified protein samples according to the instructions of the BCA protein quantitative detection kit; then performed SDS-PAGE electrophoresis, transferred membrane, blocking, and incubated with primary and secondary antibodies, ECL color rendering in order; Use imagej to perform quantitative analysis. ERK activation levels were expressed by the relative expression level of p-ERK/total ERK. All detailed statistical datasets are included in the corresponding legends. The data in the statistical chart was expressed as mean ± standard deviation, and the comparison between the two groups was performed by Student’s t-test (unpaired, two-tailed). P < 0.05 was considered statistically significant. All experiments were performed at least three times, and the results were consistent.

### Neuronal cells climbing for confocal laser imaging analysis

The coverslips were treated in advance, soaked in 1% dilute hydrochloric acid overnight, rinsed with water the next day, then autoclaved and dried, soaked in 75% ethanol for later use. The slides should be treated with 0.01 mg/ml polylysine for 10 min before inoculation, then repeatedly washed with ddH2O and dried for later transfection. The treated slides were seeded in 24-well plates one day before transfection. After 48 hours of transfection, the slides were washed with PBS, fixed with ice methanol, then stained with DAPI. Finally, the covered slides were stored in a wet and dark box for confocal laser imaging analysis.

## Results and analysis

### Introducing AzF and ONB at the TrkA-Y785 site

To introduce ONB and AzF at the TrkA-Y785 site, we constructed the TrkA-Y785amb mutant plasmid based on the obtained WT-TrkA plasmid. TrkA-Y785amb is structurally the same as WT-TrkA; that is, the TrkA intracellular domain is flagged with an EGFP tag at the C-terminus. The only difference between them is that the Y785 site has an amber stop codon mutation. So only after AzF and ONB are successfully introduced at the Y785 site, the expression of TrkA-EGFP fusion protein can be detected; that is, the expression of TrkA can be visualized by EGFP tag **(figure2a, b)**. We introduced ONB and AzF at the Y785 site in HEK293T cells by co-transfecting TrkA-Y785amb plasmid, single-plasmid system pONBYRS/U6-PyltRNA (recognizing ONB) and TrkA-Y785amb plasmid, single-plasmid system AzFRS-tRNA_4X_ (identifying AzF) in HEK293T cells. We detected the green fluorescence signals when 100 um ONB or 1 mM AzF were added to the medium, but the fluorescence was weak and undetectable without the addition of UAAs **(figure2 c,e)**.

**Figure 2.**
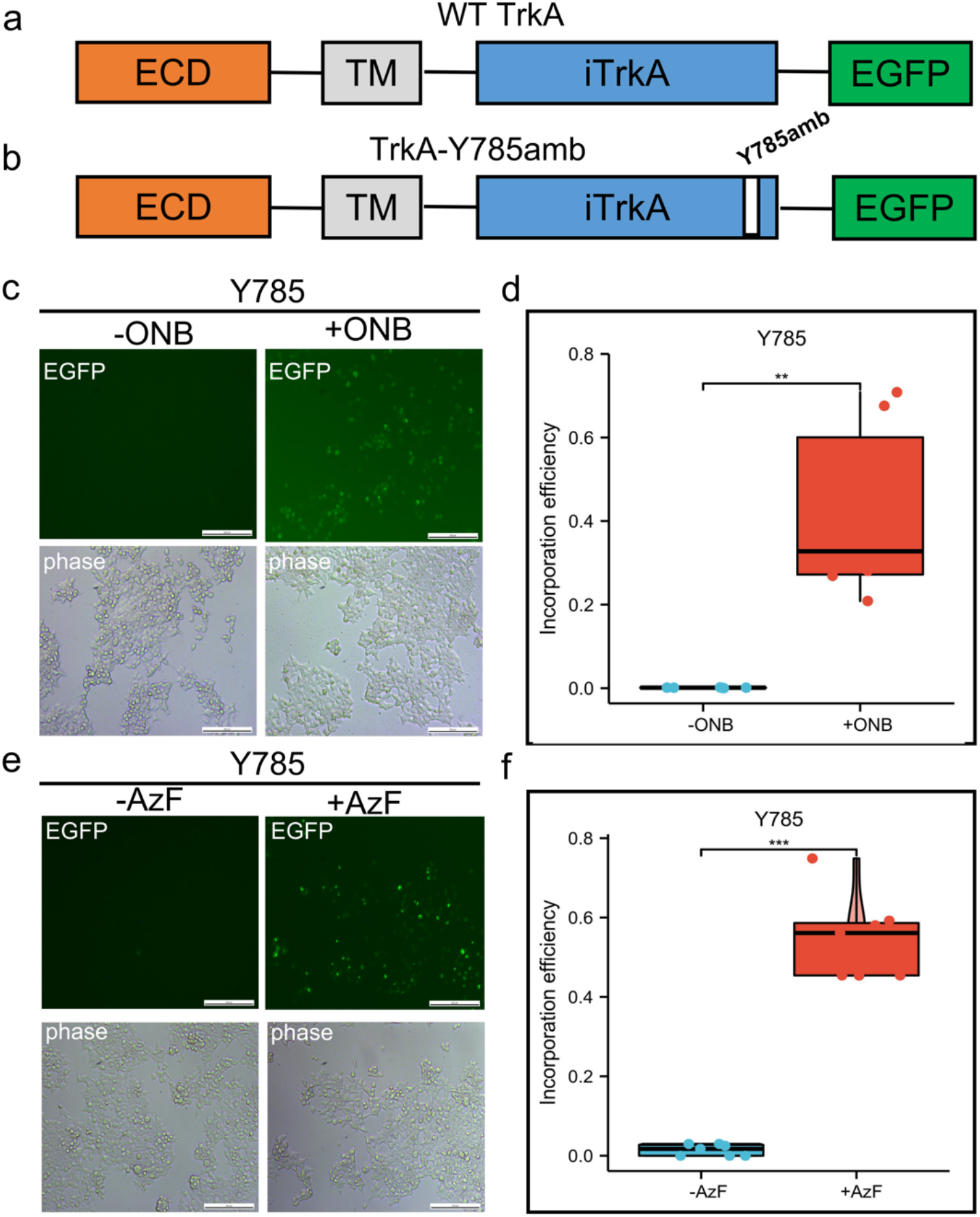
Incorporation of AzF and ONB at TrkA-Y785 in HEK293T cells. a-b) Schematic diagram of the constructed TrkA receptor tagged with EGFP at the C-term. The full-length WT-TrkA receptor includes three domains: an extracellular domain (ECD), a transmembrane domain (T.M.), an intracellular domain (iTrkA), and a fusion protein (EGFP). TrkA-Y785amb has an amber mutation TAG introduced into the Y490 position in the iTrkA domain. Fluorescent images of HEK293T cells co-transfected with AzFRS-tRNA_4X_ (AzF) or pONBYRS /U6-PyltRNA (ONB) at one site out of tyrosine kinase domain Y785 in the presence (+) or absence (-) of 100 uM ONB (c) and 1 mM AzF (e). 24-48 h post-transfection, cells were imaged using fluorescence microscopy for GFP expression (Scale bar: 200 um). (d) The incorporation efficiency of TrkA-AzF mutants in the absence (grey) or presence (dark) of AzF. (f) The incorporation efficiency expression of TrkA-ONB mutants in the absence (grey) or presence (dark) of ONB. The incorporation efficiency was described as average fluorescence intensity (%) in cells expressing UAA-incorporating receptors divided by the moderate fluorescence intensity into cells expressing the wt TrkA receptors. Mean fluorescence intensity =Indensity/area, we take a sample of six fields. Error bars show s.d.

We performed statistical analysis on the above fluorescence images. Due to a large number of fluorescent cells in HEK293T cells, we calculated the ratio of the average fluorescence intensity of Y785 to the ratio of WT-TrkA for the consideration of the difference in counts and the intensity of the fluorescence expression. The two mutants were compared with WT-TrkA(data from our previous study) to assess fluorescent expression levels and UAAs’ introduction efficiency. Analysis of fluorescence imaging results showed that the yield of TrkA-AzF mutant was about 54% of that of WT-TrkA **(figure 2f)**; in contrast, the expression level of TrkA-ONB mutant was slightly lower than that of TrkA-AzF, accounting for about 42.0% of wt TrkA **(figure 2d)**. It is indicated that the expression levels of two UAAs at the same site are slightly different. At the same time, in the absence of AzF and ONB, both groups showed quiet low fluorescence expression levels.

Thus, the difference in mean fluorescence intensity of the two mutants, as counted by fluorescence imaging, generally indicates the efficient amber stop codon suppression at the Y785 target, resulting in the specific synthesis of the AzF or ONB-containing TrkA proteins. Both tRNA/R.S. pairs exhibited good orthogonality at the Y785 site.

### Light-controlled ERK activation by TrkA-Y785 dependent on NGF treatment

In addition to the fluorescence analysis, we used the expression characteristics of the TrkA-EGFP fusion protein to reflect the read-through of TrkA. By laser confocal microscopy, we observed that TrkA-785AzF and TrkA-785ONB mutants were mainly expressed on the cell membrane, where the blue fluorescence represented the nuclear localization, and the green fluorescence represented the expression of full-length TrkA. This experiment further demonstrated that the TrkA-Y785 site could efficiently and specifically introduce AzF, ONB in HEK293T cells, thereby expressing the photosensitive TrkA-UAAs mutant on the cell membrane **(figure 3a)**.

**Figure 3.**
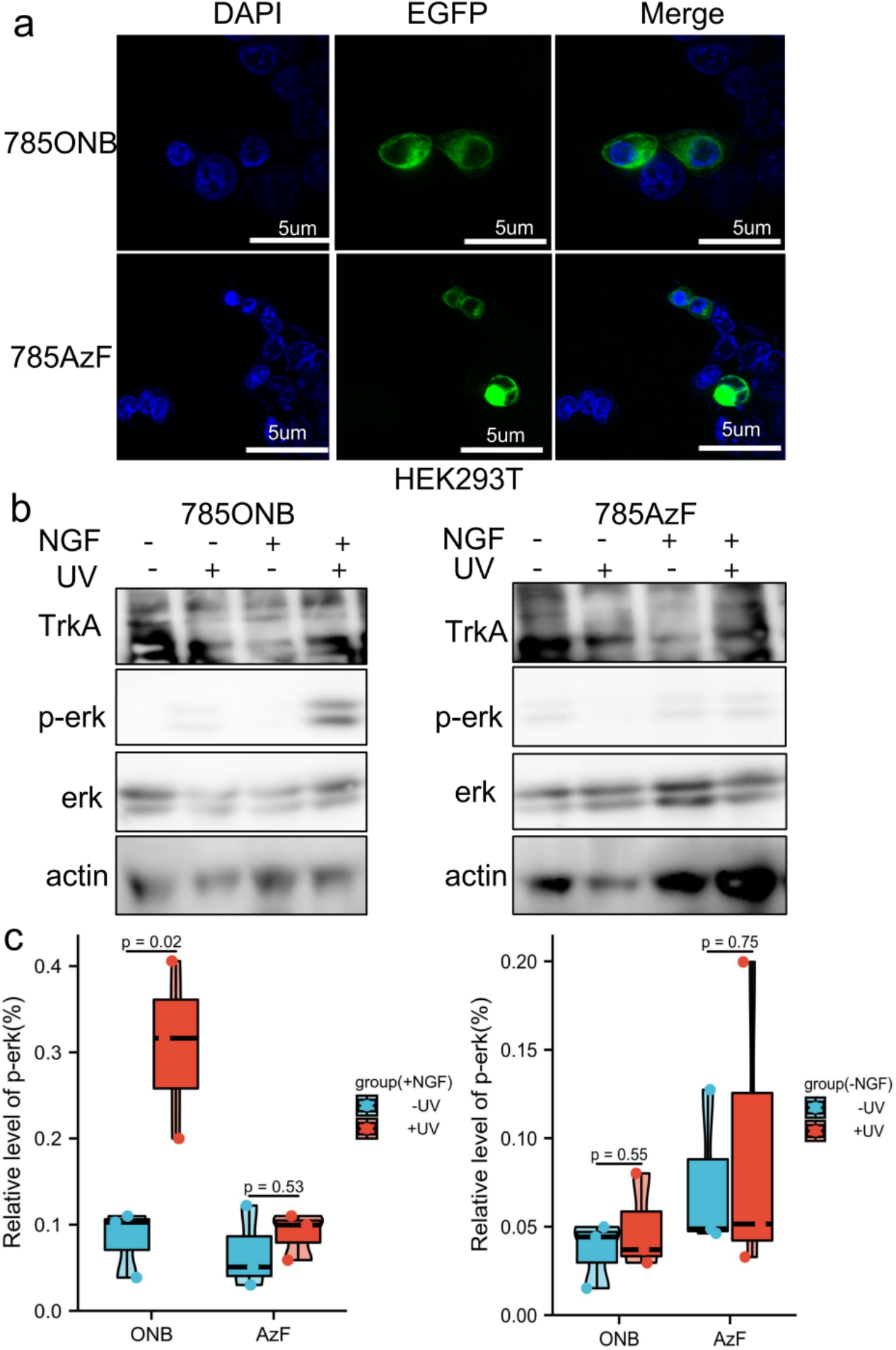
MAPK/ERK pathway activated by photosensitive TrkA-Y785ONB. a) Fluorescence images of TrkA-785AzF or TrkA-785ONB mutants expressed in HEK293T and SH-SY5Y cells. Blue, nuclei stained with DAPI (4’, 6-diamidino-2-phenylindole); green, EGFP tagged TrkA (Scale bars: 5 um). b) Western blotting analysis of HEK293T cells expressing TrkA-Y785ONB and 785AzF mutants for phosphorylated erk (Thr202 and Tyr204) and erk, without (−) or with (+) NGF and without (-) or with (+) U.V. light. TrkA and β-actin were used as a loading control. c) The expression of phosphorylated erk for TrkA-785ONB and 785AzF mutants in the absence (blue) or presence (red) of U.V. The cells were transfected with amber mutants at the Y785 site of TrkA and R.S./tRNA pairs for each UAA, expressing TrkA-ONB and TrkA-AzF at Y785 in the presence and absence of NGF. Error bars show s.d. p<0.05 represents statistically significant.

Next, to test whether this light-controlled strategy testing at the Y785 site could activate the MAPK/ERK pathway, we examined the activation of p-ERK by TrkA-Y785ONB and TrkA-Y785AzF mutants after NGF and U.V. treatment. Under U.V. light irradiation, ONB released the “photocage” group from the introduction site and recovered to tyrosine, followed by autophosphorylation of the hydroxyl group on the tyrosine site. To assess whether photoactivation of TrkA-Y785ONB leads to functional changes in ERK signaling, we performed immunoblot analysis of p-ERK activity following U.V. treatment using live cells which express photosensitive UAAs. The TrkA-Y785ONB mutant was expressed in HEK293T cells by co-transfection of Y785amb plasmid and pONBYRS/U6-PyltRNA pairs. Then cells were treated with NGF and U.V. stimulation, and then extracted protein and detect TrkA, p-ERK, ERK and actin levels. It is found that p-ERK was not detected in TrkA-Y785ONB in the absence of U.V. light irradiation due to the presence of photolabile groups in ONB, while the U.V. stimulation-induced apparent expression of p-ERK **(figure 3b)**. Quantification of p-ERK levels revealed that TrkA-Y785ONB exhibited an expression level of about 30% under U.V. stimulation and showed a stronger expression level than the untreated U.V. group (8.3%) **(figure 3c)**. We also examined the photo-induced effect of AzF at the Y785 site. AzF undergoes a different photochemical process compared with ONB. Upon U.V. treatment, AzF typically generates a nitrogen radical, which may subsequently be reduced to an amine or form a covalent bond with an adjacent atom within a 3–4 Å distance. Unlike the previous results in the kinase domain region^**14**^, the Y785 site did not show an apparent photo-controllable activation level of p-ERK after the introduction of AzF. However, in the absence of NGF stimulation, there was no significant difference in p-ERK expression levels with or without light in two mutants **(figure 3b)**.

Taken together, our results demonstrate that TrkA-Y785 can induce p-ERK activation by light control only in the presence of NGF. Among them, ONB acts as a powerful light-controlled “switch” to control the phosphorylation state of this tyrosine site.

### The photosensitive TrkA-Y785ONB mutant regulates SH-SY5Y cells differentiation dependent on NGF

Compared to HEK293T cells, which were able to efficiently transfect and successfully express TrkA-ONB, TrkA-AzF mutants, the successful application of this technology to neurons required extensive experimental optimization. We have tested light-sensitive TrkA mutants (Y490, Y670, Y674, Y675) in SH-SY5Y cells in our previous study to establish a light-controlled method for neurite outgrowth^14^. Therefore, using the same strategy, we introduced AzF and ONB at the Y785 site and carried out TrkA-AzF and TrkA-ONB fluorescence expression detection in SH-SY5Y and PC12 cells. In the case of adding AzF or ONB in the medium, only weaker EGFP expression could be detected in two groups of cells, but the number of green fluorescence cells in PC12 was weaker than SH-SY5Y cells **(figure 4a).** In the absence of AzF or ONB, almost no fluorescence was detected. Fluorescence statistics of EGFP expression levels in two cells showed that the average yield of TrkA-785AzF mutant in PC12 cells was about 2%, while TrkA-785ONB was less than 1%, showing a low introduction efficiency. In SH-SY5Y cells, the average yield of TrkA-785AzF and TrkA-785ONB mutants was about 7%, and the average expression levels of the two UAAs were similar **(figure 4b)**. In addition, compared with PC12 cells, SH-SY5Y cells exhibited higher incorporation efficiency of photosensitive UAAs at the TrkA-Y785 site.

**Figure 4.**
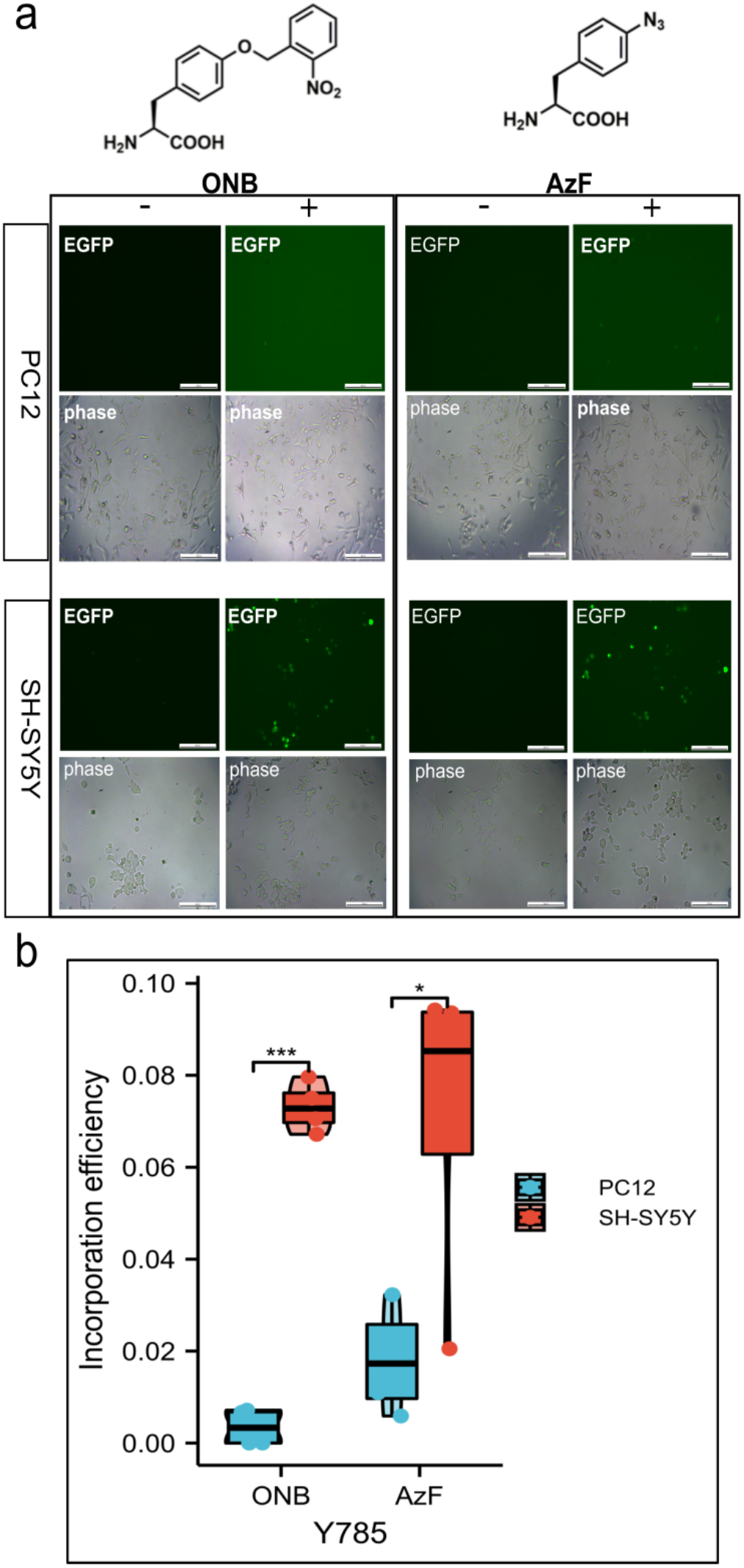
The incorporation efficiency of ONB at Y785 sites. a) fluorescent images of HEK293T cells and SH-SY5Y cells co-transfected with AzFRS-tRNA_4X_ (AzF) or pONBYRS /U6-PyltRNA (ONB) at one site out of tyrosine kinase domain Y785 in the presence (+) or absence (−) of 100 uM ONB (left) and 1 mM AzF (right). 24-48 h post-transfection, cells were imaged using fluorescence microscopy for GFP expression (Scale bar: 200 um). b) Average expression of TrkA-AzF, TrkA-ONB mutants measured for indicated conditions in the presence of 1 mM AzF or 100 um ONB, quantified by the percentage of fluorescent cells (EGFP) to total cells (phase). Error bars show s.d.

In addition, immunoblotting experiments showed that in WT SH-SY5Y cells with or without NGF treatment, the p-ERK signal was almost undetectable **(figure 5a)**. After transfecting exogenous WT TrkA in cells, the ERK activation could only be detected after NGF treatment **(figure 5a)**. We tested SH-SY5Y cells expressing WT TrkA and found that after treatment with 50 ng/ml NGF for 24-48 h, cells differentiated significantly with or without U.V. light treatment. No obvious cell differentiation effect was seen while cells were treated without NGF stimulation. **(figure 5b)**,which indicated that the exogenous TrkA retained ligand sensitivity and signaling activity, and U.V. light had no obvious effect on cell differentiation.

**Figure 5.**
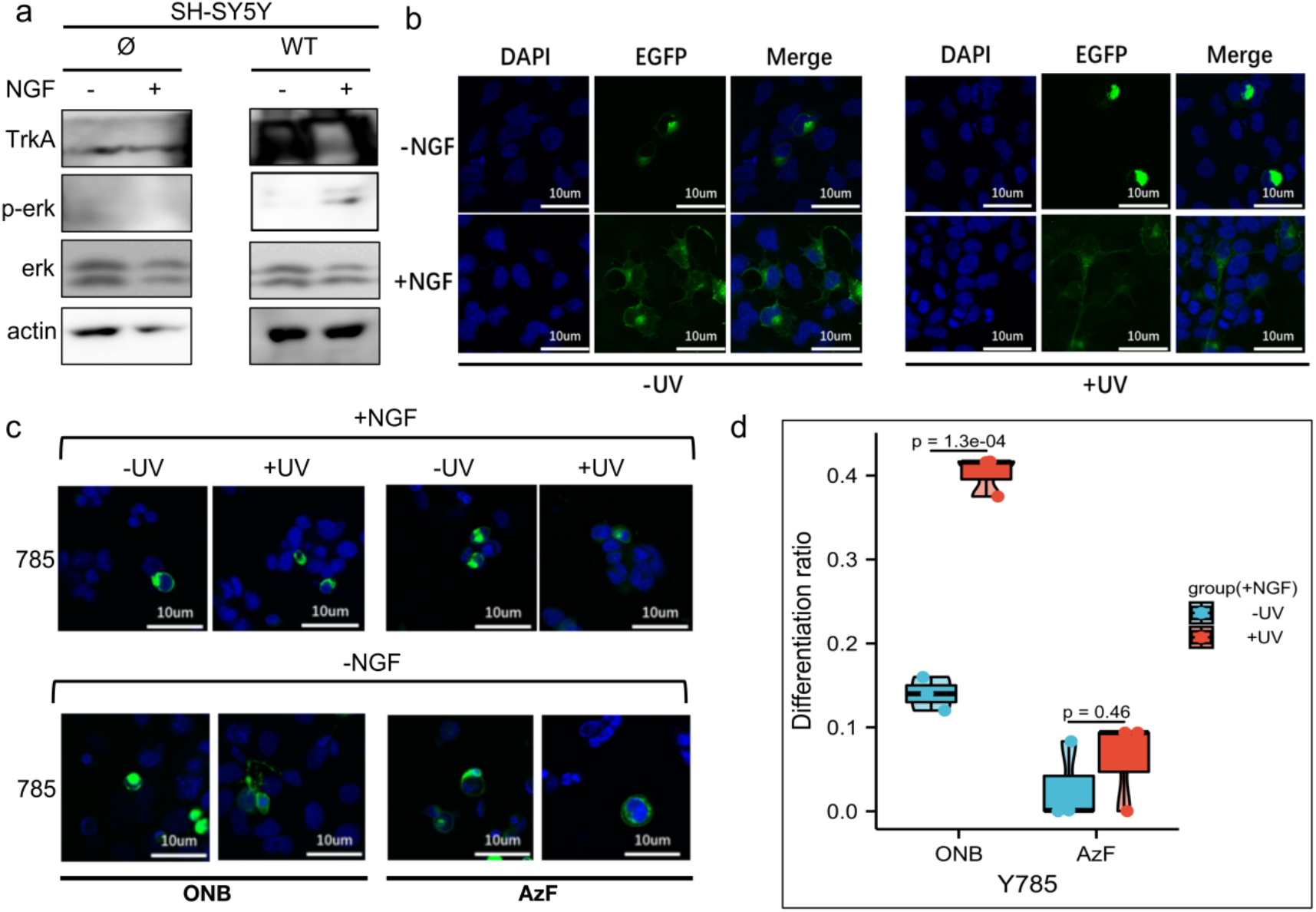
TrkA-785ONB mutant promotes SH-SY5Y cell differentiation in response to U.V. stimulation with NGF. a) Western blotting analysis of SH-SY5Y cells expressing endogenous TrkA (Ø) or wt TrkA for phosphorylated erk (Thr202 and Tyr204) and erk without (−) or with (+) NGF. TrkA and β-actin were used as a loading control. (b) Representative fluorescence images depicting SH-SY5Y cells expressing wt TrkA before (−) and after (+) U.V. treatment, with medium supplied without (−) or with (+) 50 ng/ml NGF kept in the dark for 24-48 h before imaging. c) Representative fluorescence images depicting SH-SY5Y cells expressing TrkA-785ONB (two columns on the left) or 785AzF (two columns on the right) mutants incorporated at indicated sites with (+) and without (−) U.V. treatment, with medium supplied with NGF (50 ng/ml). Cells were cultured in the cell incubator for 24-48 h (dark) before imaging. d) Differentiation ratios calculated for various mutants expressed in SH-SY5Y cells in the presence of NGF with or without U.V. light. Differentiation ratios were calculated as the following: taking the ratios (No. of green fluorescing cells with neurite longer than cell body/No. of green fluorescing cells) for Y785 and wt TrkA at each condition. Then take the ratios of each mutant/wt TrkA treated in the same condition. Values above represent the mean ± S.D. of three biological replicates (n = 3) with >200 cells counted per replicate. Scale bars:10 um.

We have successfully expressed light-sensitive UAA-mutants at the TrkA-Y785 site in SH-SY5Y cells. The next step was to analyze neurite outgrowth and cell differentiation with and without NGF treatment through the light-controlled Y785 site. TrkA-785ONB and TrkA-785AzF mutants were expressed by co-transfecting this set of UAA coding systems into SH-SY5Y cells, which were consistent with the previous western blotting treatment. Neuronal cells were cultured in the dark for 24-48 h in the serum-free medium containing 50 ng/ml NGF, and their neuronal growth was compared with WT TrkA. In the presence of NGF, cells expressing the Y785ONB mutant showed differences in UV-induced differentiation relative to the untreated U.V group (differentiation rates: UV-treated group: 40.6%; UV-untreated group: 13.9%) **(figure5c, d)**. However, the mutant Y785AzF without obvious ERK activation effect before and after U.V. irradiation showed no obvious difference in cell differentiation before and after U.V. treatment (differentiation rates: UV-treated group: 0.6%; UV-untreated group: 0.3%). In the absence of NGF treatment, we observed no apparent cell differentiation phenotype for either UAAs **(figure 5c)**.

Thus, our results suggest that activation of p-ERK in the TrkA-Y785 site exhibits substantial consistency with neuronal cell differentiation regardless of NGF stimulation.

## Discussion

The general concept of TrkA signaling activation is that the ligand NGF from the extracellular domain promotes the homodimerization of TrkA at the cell membrane, which in turn activates downstream signaling pathways. However, because the TrkA signaling pathway is intricate and related to five intracellular phosphorylation sites (Y490, Y670, Y674, Y675, Y785) and their combinations, the mechanism has not yet been fully elucidated^19^.

We have been interested in developing systems for light-controlled protein function through site-specific control of the kinase receptor TrkA. Using Genetic Code Expansion (GCE)^20,21,22^ technology to combine photosensitive UAAs with site-directed mutagenesis (phosphorylation sites) for precise, light-directed activation of the membrane kinase receptor TrkA, this method provides a novel concept for studying NGF/TrkA signaling pathways. At present, we have tried at five phosphorylation sites of TrkA (two docking sites Y490, Y785, and three sites Y670, Y674, Y675 in the kinase domain) and found that the downstream MAPK/ERK pathway can be specifically activated by light-controlled means through the introduction of photosensitive UAAs into these five sites. The two introduced molecules, ONB and AzF, are chemically structurally distinct from tyrosine and both can prevent phosphorylation before photoactivation. However, different photochemical reactions will occur after U.V. illumination^23,24,25^, which provides a light-controlled method for analyzing the contribution of each tyrosine to the activation of specific pathways during TrkA signal transduction. This advantage cannot be provided by conventional mutagenesis methods^26,27^.

Our light control study of the TrkA-Y785 site indicated two points of interest: 1) The difference of the Y785 site on ONB and AzF. Ligand-dependent activation of Y785ONB may reflect a ligand-induced site-state-dependent activation process; 2) The two docking sites, Y490 and Y785, are the main activation sites to MAPK/ERK pathway. SH-SY5Y cells were significantly differentiated at the Y785 site compared with before and after light control under the condition of introducing ONB and NGF treatment. Light-controlled activation of the Y785 site indicated that this site activates the MAPK/ERK pathway dependent on the NGF ligand. And using the SH-SY5Y cell model with less endogenous TrkA interference, we combined this detection of the ERK pathway (microscopic) in photosensitive TrkA mutants with neural cell differentiation (cellular phenotype) and found they all showed good consistency. This is reasonable because Y785 has been reported in previous studies to activate ERK in a PLCγ-PKC signaling-dependent manner, and cell differentiation was significantly reduced after simultaneous inhibition of both Y490 and Y785 phosphorylation, suggesting that Y785 makes important contributions in regulating the ERK pathway. Our light-controlled approach offers advantages in explaining the phenotypic roles of the Y785 site and other single tyrosine sites^14^. We expect that through separation and combination light-control studies of these critical sites, it will provide a powerful approach to resolve the polymorphism of tyrosine receptor phosphorylation and the diversity of its functions.

To sum up, we supplemented light control results of the TrkA-Y785 site based on our previous research and further expanded the application of genetic code expansion technology in kinase receptors; Combined with the previous research results^14^, we propose that these five phosphorylation sites may have different contributions to the activation of the MAPK pathway, and site-dependent phosphorylation in TrkA may have polymorphic control over the signaling process. Through precise optical control^28–32^, the activity of kinase receptors can be regulated at a single phosphorylation site to achieve the effect of site-directed light-controlled phosphorylation. This approach lays the foundation for comprehensive analysis of kinase-related pathways and screening of compounds that intervene in site-specific phosphorylation pathways in targeted therapy.

