## Supplemental Figure for "Light-controlled phosphorylation in the TrkA-Y785 site by photosensitive UAAs activates the MAPK/ERK signaling pathway"

**Supplmentary figures**

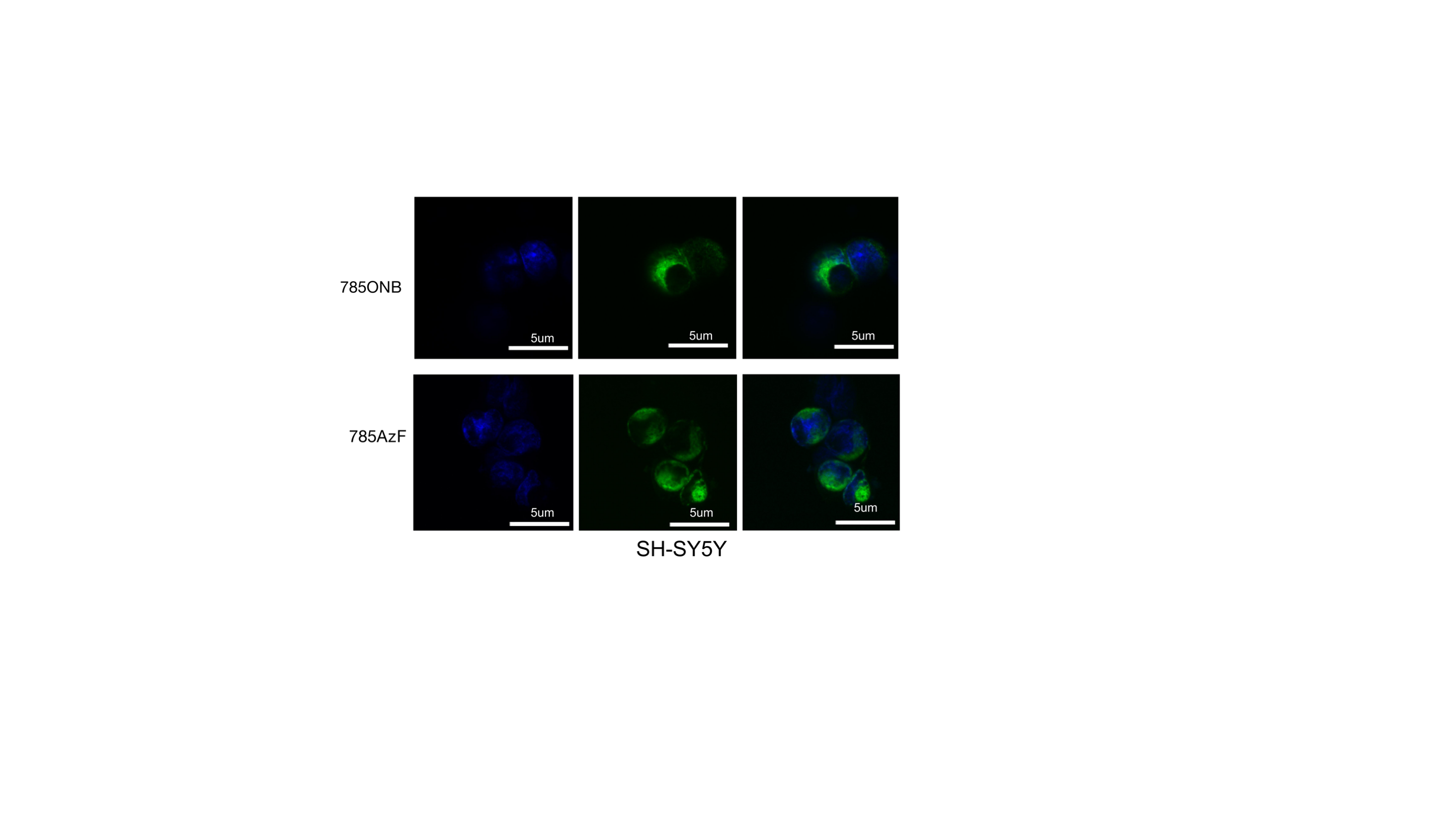

**Figure 1 Localization of TrkA constructs in SH-SY5Y cells** Fluorescence images of wt TrkA receptors and TrkA-785AzF or TrkA-785ONB mutants expressed in SH-SY5Y cells. Blue, nuclei stained with DAPI (4’, 6-diamidino-2-phenylindole); green, EGFP tagged TrkA. (Scale bars: 10 um)

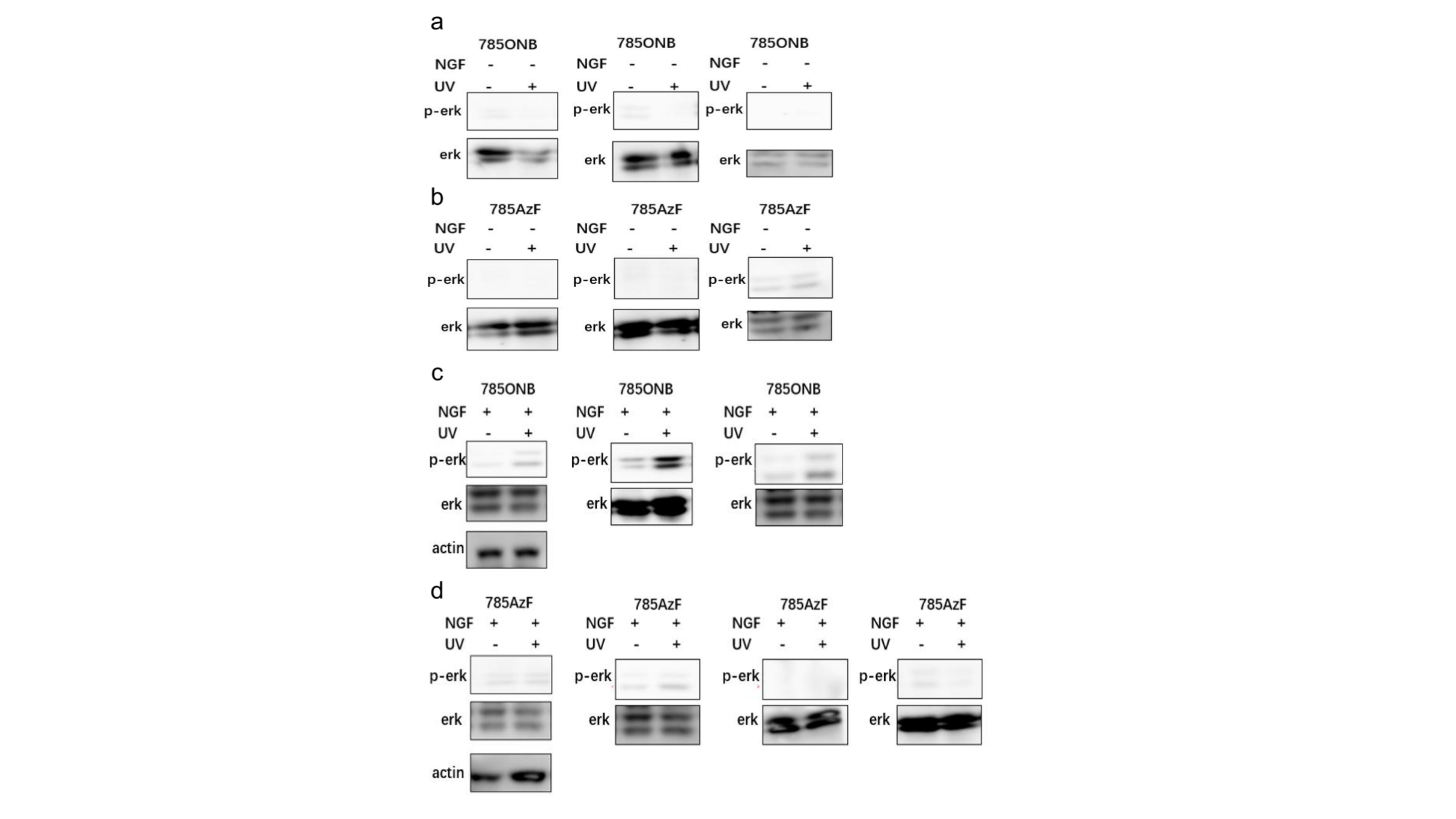

**Figure2 Repeats for gel experiments** Western blotting analysis of HEK293T cells expressing TrkA-785ONB and TrkA-785AzF mutants for phosphorylated erk (Thr202 and Tyr204) in the presence or absence of ligand NGF without (-) or with (+) UV light.
